# Thin Film Metallization Stacks Serve as Reliable Conductors on Ceramic-based Substrates for Active Implants

**DOI:** 10.1101/2020.06.25.171454

**Authors:** Patrick Kiele, Paul Čvančara, Michael Langenmair, Matthias Mueller, Thomas Stieglitz

## Abstract

Hermetic and non-hermetic packages of active implantable medical devices are often fabricated of ceramics like alumina. Screen printed PtAu paste is the state of the art metallization for functional structures. Due to solid state and liquid diffusion of Au at thermal exposure, solder times are limited; otherwise metal structures tend to delaminate. Moreover, it was shown that PtAu with solder fails after 37.4 years. We established a thin film metallization on alumina process to overcome these disadvantages. The metallization consists of sputtered platinum with an underlying adhesion layer made of tungsten-titanium to increase the adhesion strength to the alumina substrate. We avoided using gold in this work due to its high diffusion tendency. Instead, the materials in use provide relatively low diffusion properties, which potentially increases the long term mechanical performance and usability during assembly and packaging.

Utilizing the Design of Experiment (DoE) methodology, we derived an optimal Pt thickness of 500 nm with 43 nm of WTi as adhesion promoting layer. After accelerated aging at 150 °C, corresponding to 125 years at body temperature (37 °C), the contact pad adhesion strength was with 32.75 MPa ± 7.08 MPa still significantly higher than the safety limit of 17 MPa, following the recommendations for a robust screen-printing metallization process. Moreover, soldering times of up to 120 s did not influence the adhesive strength. The new process reduced the minimum track distance to 50% of screen printing values and is capable of rapid prototyping. It helps to make the assembly process independent of the manufacturing person, to increase the yield of device fabrication and -most important in implantable device manufacturing-to make it more robust and thereby more safe for the patient.

## I. Introduction

**C**eramics like alumina are widely used within the medical industry. The applications of alumina range from dental implants [1] over femoral heads [2] to bioinspired functional material composites [3], just to mention a few. However, this work focus on the usage of ceramic materials in the manufacturing of active implantable medical devices (AIMDs) like brain computer interfaces (BCI) for recording and stimulation of brain tissue [4], retinal implants for restoration of sights [5, 6] or cochlear implants for stimulation of the auditory nerve [7]. In these applications alumina is used widely as a substrate material for electrical circuits, electrical insulators or as a hermetic shell to protect the implant electronics from the surrounding tissue and vice versa. Ceramics exhibit advantages like mechanical strength, high hermeticity, are electrochemically stable and therefore do not corrode within the body [8]. Ceramics can also be used within non-hermetic packaging as substrate for printed circuit boards (PCB) to interconnect, for example, polymer based thin film electrodes with transcutaneous cables [9]. Contrary to epoxy based PCBs, ceramics have nearly no water uptake, which would lead to delamination of the functional layers. Additionally, ceramics show superior properties in telemetry in contrast to metals as they are permeable for electromagnetic waves and less imaging artefacts in magnetic resonance imaging (MRI) [10]. Therefore, the antenna can be implemented within the housing, reducing the number of feedthroughs. In the application field of AIMDs the electronic system including the ceramic parts is commonly casted in silicone rubber to ensure biocompatibility to the surrounding soft-tissue and simultaneously protecting exposed metallic components from corrosion [11]. Therefore, the excellent adhesion properties to silicone rubber are crucial for both non-hermetic and hermetic packaging to avoid osmosis due to gaps or voids and therefore delamination and eventual failure of the silicone rubber encapsulation. [12]

PCBs can contain electrical circuits as well as feedthroughs and interconnection structures. These PCBs for AIMDs are fabricated using a ceramic as substrate and depositing the desired functional thick film structure via screen-printing [13, 14]. The track distance is limited to about 0.125 mm [15] due to the minimal mesh size resulting in inaccurate edges. The screen printing paste consists of a glass frit, an organic carrier and the functional material, e. g. PtAu. By firing the paste, the organic carrier is removed, the functional material melts and interconnects within the glass frit, which in turn builds up covalent bonds towards the ceramic. The drawbacks of this method are, that the functional material exhibits only mechanical and low force bonds like van der Waals forces and that thermal mismatch of the components can induce residual stresses after firing [16, 17]. Kohler [18] showed, that PtAu as screen-printing paste exhibits disadvantages due to solid-state and liquid diffusion processes while fabrication and within application. Thermal exposure like soldering at 350 °C leads within 20 seconds to entire diffusion of gold into the solder and as a consequence to delamination of the metallization. In case of materials difficult to solder like cobalt base alloys, this process increases the risk of implant failure. Moreover, it was shown with accelerated aging that failure between substrate and metallization occurs after 37.4 years at body temperature (37 °C). For chronic applications like cochlear implants which should last for a lifetime of about 80 years, this would be equivalent to a total failure.

This study was designed to overcome the issues of the described thick film screen printing metallization. In order to meet the constraints of implantable medical devices, especially to avoid toxic additives that could be present in electroplating processes, our strategy was to use sputtered slow diffusion materials only to increase the long-term reliability and to allow time consuming assembly processes like soldering. Therefore, we avoided gold in the metallization layer and used pure platinum only. Further, the initial bonding strength of the platinum metallization to the alumina substrate was increased by adding an adhesion layer (WTi), which can provide chemical bonding mechanisms to both alumina and platinum. By changing the fabrication process to a sputtering process, the chemical bonding mechanisms were supported. High energy processes like sputter deposition of metal thin films can lead to an atom implantation into the ceramic lattice and in parallel to surface diffusion, whereby covalent bonds between the ceramic and thin film metallization are formed, on condition that the materials are chemically compatible.

Functional structures like tracks and solder pads are realized by removal of excessive material with laser ablation, with further advantage of rapid prototyping capability. The minimum track distance is defined by the diameter of the laser spot (in our case 54 µm [19]). Furthermore, the structuring can be easily translated to commercial large-scale microelectronics manufacturing technologies like lift-off processes to allow mass fabrication for the final product. Our hypothesis is, with using physical deposition methods like sputter deposition of platinum and an adequate adhesion promoter, the bonding to alumina is dominated by covalent bonds instead of physisorption and mechanical interlocking. This increases process stability, eliminates the influence of solder times and improves structural precision. Moreover, it increases the long-term-stability, which would lead to less risk and higher safety for the patient. In addition, the metallization process parameters are optimized with respect to the initial adhesion strength to the ceramics and the electrical conductivity by performing the engineering approach of a Design of Experiment (DoE) rather than a FEM based evaluation. The collected data in combination with a DoE is more promising to reflect the process reality in our study due to the interactions of different bonding mechanism and boundary conditions

## II. Fundamentals of Adhesion

The following chapter gives a brief overview of the fundamentals of the relevant adhesion mechanism theories used in this work. For further reading we recommend the study of specialized literature on adhesion, e.g., [20].

Adhesion in general is the ability of two different materials to stick together. However, the adhesive process is a very complex field, where several adhesion theories have been described in literature [21]. A common consent is that adhesion emerges of an interplay of the different adhesion mechanisms described below: mechanical interlocking and adsorption [22].

Surface roughness allows *a mechanical interlocking* of both phases. The interlocking originates in (micro) pores and capillaries of the surface [23]. Regarding the sputter deposition, single atoms or clusters solidify in the pores and interlock with the substrate while the complete layer is formed. Nevertheless, (micro) pores and capillaries increases the active surface providing room for adsorption mechanisms, which play a major role in the theory of adhesion [20]. Accordingly, the surface roughness of the substrate (i.e. the ratio between the active and the geometrical surface) correlates with the adhesive forces.

*Adsorption* can be divided depending on the bonding species in physisorption and chemisorption. Physisorption accrues based on intermolecular attraction [22], with electrostatic van der Waals bonds occurring due to permanent and or inducted dipoles [24]. This adhesive force decreases with the sixth power of the distance between the centers of gravities. Therefore, physisorption is effective in the range of two inter-atomic distances, only [25]. However, chemisorption is based on chemical bonds such as covalent, ionic and hydrogen bonds (50-500 kJ/mol or 0.52-5.18 eV) [24, 26], which results in stronger bonds than the physisorption (<50 kJ/mol or <0.52 eV) [26]. While physisorption occurs with any kind of atoms, chemisorption requires compatible materials that can react with each other [27].

## III. Materials And Methods

### A. Surface metallization and adhesion promoter

Platinum provides great potentials for implantable medical devices. Despite of its relatively high price and lower conductivity compared to copper, gold and silver, which makes this material a rare choice in the common manufacturing of PCBs, the inert properties of platinum prevent surface oxides to be developed at standard conditions[28] as well as in oxidizing atmospheres of up to 800 °C [29]. Further chemical or mechanical processing are not necessary for soldering and bonding [28]. Beside this, platinum features excellent biocompatibility and low diffusion rates into e.g. SnPb based solder [28], two important factors concerning the manufacturing and long term stability of implantable devices. The nature of the substrate and the platinum metallization do not allow direct chemical interactions. The reason for this is that the metallic bonds of platinum are not compatible with the bonds in alumina, which are of covalent nature with ionic fractions [22]. Therefore, the adhesion of platinum on a ceramic substrate is rather of physical than of chemical nature [22, 30] and has to be improved in order to deliver a stable performance for implantable devices in medical applications [31]. To circumvent these issues and increase the adhesion, it is possible to integrate an adhesion promoter between the ceramic substrate and the functional platinum metallization [22] (e.g. titanium [29, 31, 32], platinum oxide [31–33]).

Sputter-deposited titanium can serve as an excellent adhesion promoter on ceramic substrates to sputter-deposited platinum. Critical for this application is the increase in adhesion and the stability of the material stack. In regard to the first criterion, the affinity of titanium to form oxides potentially enables it to chemically bond to the ceramic substrate (Al_2_O_3_) [27]. During sputtering, titanium atoms are introduced in the substrate’s lattice structure, forming a first unstable nucleus. Within the ongoing sputter deposition process the nuclei are growing and forming a bigger sized chemisorbed stable cluster. [34] In this scenario, titanium can form a covalent bond with the oxygen atom of the ceramic substrate which leading to a mixed oxide (Al-O-Ti) [22], which is essential for interfacing ceramic substrates [35]. Mechanical interlocking and physisorption can be considered to only play minor roles in this case due to their smaller strengths.

Regarding the whole material stack, titanium features stable performance with low diffusion into the adjacent platinum at temperatures up to 200 °C [29]. Consequently, care has to be taken also during fabrication to keep the substrate temperature at moderate levels during sputtering-deposition of the platinum layer[31]. Another advantage of titanium as adhesion promoter to platinum is the reliable performance as a pad material for soft soldering (via chemisorption [28]) as well as for gold wire-bonding (via physisorption [36]). This (in contrast to the previously investigated platinum oxide [32]) renders titanium to be suited for the proposed application in wet environments at 37 °C. Even though many other currentless electrodeposition technologies exist and are applied in microelectronics and sensors R&D, we omitted these intentionally due to their use of potentially toxic chemicals (e.g. dimethylamine borane-DMAP) and metals (e.g. copper). The use of these materials is strongly counter indicated for implantable devices.

### B. Fabrication process & sample preparation

Cleaning: To achieve proper adhesion, cleaning of the substrate prior to the deposition of the metallization is required[18]. We used cleaning procedure that is used in different groups in ceramic based medical implant development consisting of shaking in “Leslie’s soup” (Teepol-L, Teepol Products, Kent, UK, mixed with sodium phosphate and DI-water) (5 min) followed by rinsing with isopropanol and DI-water [37–39]. The substrate was finally treated in an ultrasonic bath (10 min) and dried with pressured air.

Metallization: The thin film metallization consisted of a layer stack made of sputtered tungsten-titanium forming an adhesion layer and platinum (Leybold Univex 500, Leybold Vacuum GmbH, Cologne, Germany) on a sintered alumina (Al2O3, 96 %) substrate. Alumina was chosen as it provides excellent mechanical properties and good biocompatibility [40]. Furthermore, it provides a high diffusion barrier against water, which prevents a swelling of the substrate and therefore intrinsic stress at the metal interface [14].

Structuring: In this work, we decided to structure our samples with a subtractive laser process that allows high flexibility in the design phase. The material removal was done by thermal evaporation utilizing a high-energy nanosecond laser (DPL Genesis Marker, ACI Laser-Components, Nohra, Germany) with a wavelength of λ = 1064 nm (Figure 1B).

**Figure 1:**
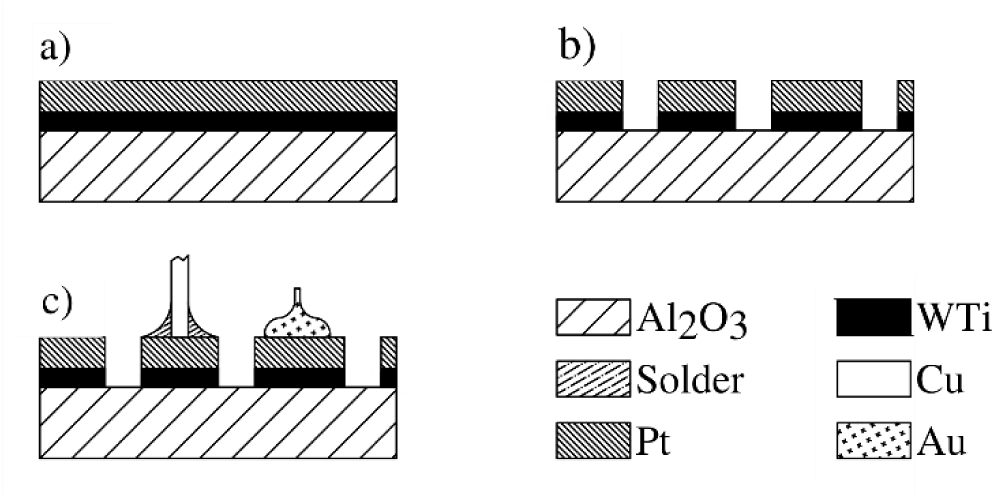
Fabrication process of samples used to characterize the Al2O3-WTi-Pt layer stack under investigation. A) Sputtering of an adhesion promoter (WTi) followed by the surface metallization (Pt) B) Laser structuring of the metallization layers C) Assembly of solder joints and gold ball-bond studs for characterization.

Assembly: Electrical contact was established by soldering and bonding processes. Soldering was done manually with a solder iron (Tsolder = 350 °C) on top of a hot plate (Tplate = 120 °C) using a lead based solder (Sn62Pb36Ag2). Bond studs (e.g. to establish Microflex-interconnections[41]) were applied using a 25 µm thick gold-wire in a standard wire bonder (K&S 4524, Kulicke & Soffa Pte Ltd, Singapur) (Figure 1C).

### C. Optimization of processing parameters

As there is a huge variety of possible process parameters, a Design of Experiment (DoE) was performed to develop a mathematical forecast model that reflects the correlation of the chosen input parameters and the investigated output values. The DoE was executed using the software JMP (v. 12.2.0, SAS Institute Inc., Cary, North Carolina). Important input parameter sweeps were performed on the thicknesses of the adhesion layers (*z*_*WTi*_) and surface metallization (*z*_*Pt*_) as well as the annealing temperature *T*_*anneal*_ (*t*_*hold*_ = 4 h) [33]. Additionally, the time of soldering (*t*_*solder*_) was varied while keeping the soldering temperature constant at 350 °C to evaluate the independency of the user’s experience in soldering [18]. The four parameters and their investigated factor level combinations (response surface method, central composited design) are listed in table 1. Each factor level combination was measured ten times (n = 10). The output parameter of the soldered connections was represented by the adhesive strength (σ_*M*_) of the layer stack, whereas the output parameter for the gold studs was set by the maximum shear force.

**Table 1:** Investigated parameters with their minimum (-), center (o) and maximum (+) factor level. In accordance to this 4x3 parameter matrix, 16 configurations got manufactured and tested. Additionally, one set of parameters was used to fabricate samples for accelerated aging.

| Parameter | Variable | Factor level combinations |  |  | Process parameters - accelerated aging |
| --- | --- | --- | --- | --- | --- |
|  |  | - | o | + |  |
| Thick. adh. Layer (nm) | $z_{WTi}$ | 10 | 55 | 100 | 43 |
| Thick. Pt-layer (nm) | $z_{Pt}$ | 100 | 300 | 500 | 500 |
| Annealing temp. (°C) | $T_{anneal}$ | 20 | 360 | 700 | - |
| Time of soldering (s) | $t_{solder}$ | 2 | 11 | 20 | 2-5 |

### D. Layer characterization

In order to characterize the properties of the above described surface metallization on the Alumina, scanning electron microscopy (SEM) in combination with focused ion beam (FIB, Zeiss Auriga 60, Carl Zeiss AG, Oberkochen, Germany) was utilized. With the FIB it is possible to remove a defined amount of material and to prepare the resulting side walls with a high surface quality. Using the SEM, high resolution cross-sections of the thin film metallization and the underlying ceramic could be acquired. The investigated sample was fabricated according to section 3.2 and placed on the conventional SEM sample holder.

### E. Mechanical performance

The mechanical characteristics and stability can be quantified with a broad range of test designs. Most important for the application on hand is the tensile strength of the investigated material stack. A multitude of methods testing for maximum tensile stress exist. Modern scientific opinion suggests that torsional loading until failure produces the most representative material characteristics [20, 42].

Introducing innovative materials in the medical industry is only possible when new approaches can exceed the performance of state of the art technology. Therefore, this work is evaluating the novel material stack against safety limits stated by material suppliers of traditional glass-based pastes and is comparing the results to excessive reliability studies published in the medical field [18]. These studies utilize axially loaded butt joint tests. Since the comparability of maximum tensile strength values are highly dependent on the test setup [20, 43] a variation of butt joint loading is the method of choice for the solder joined connections in this work [16].

The test setup for the solder joined connections contained 15 mm (Ø = 0.8 mm) copper (Cu) wires manually soldered onto 1 × 1 mm^2^ solder pads (Figure 2a) that were fabricated using the described method with their respective process parameter combinations (Table 1). The prepared samples were fixated in a blocking frame (Figure 3) and pull tested until destruction recording the applied force and the dislocation of the pulling clamp (Dage4000, Nordson GmbH, Erkath, Germany; Cartridge: WP10kg) at a pull speed of 500 µm/s. Being a variation to traditional butt joint pull tests the applied holding structure assures the pulling force to be applied locally at the areas of interest, while preventing the brittle base material (the alumina substrate) to bend and break.

**Figure 2:**
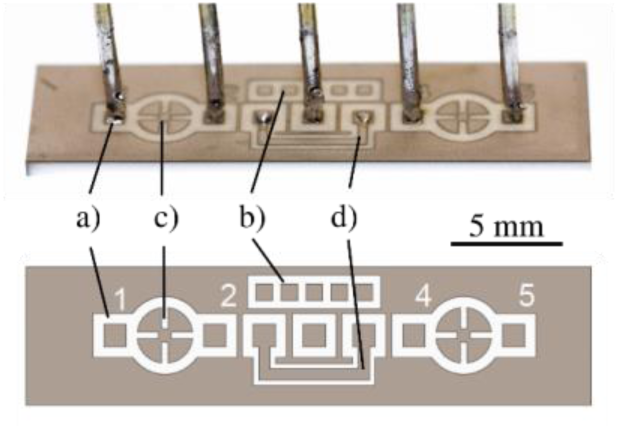
Schematics and assembled test structures for a) pull testing, b) shear testing of ball studs, c) Van-der-Pauw measurements and d) el. contact evaluation of solder joints.

**Figure 3:**
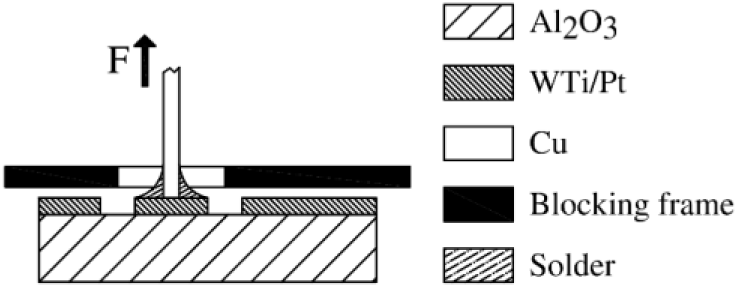
Configuration for adhesion testing of the thin film metallization using a pull test including a blocking frame to prevent damage of the ceramic substrate.

The material stack of the gold stud bonding process was evaluated with a shear test setup (Dage4000, Nordson GmbH, Erkath, Germany; Cartridge: BS5Kg; Chisel width: 300 µm) at a constant shear height (5 µm) and speed (250 µm/s) as proposed in other publications [36, 36, 44, 45]. The bond studs were fabricated on 0.65 × 0.65 mm^2^ metal pads (Figure 2b). The shear force was traced and the type of breakage (cohesive failure in the gold stud, adhesive failure, mixed failure) was graded.

Based on the optimized parameters resulting from the performed DoE, the long term performance of the mechanical integrity was evaluated. Test structures were designed and developed (Figure 2) to investigate the adhesion strength of the choses metal stacks to the ceramics after soldering wires and stud-ball bonding.

#### Alteration during soldering

Further evaluation of the influence of liquidus diffusion during soldering was further observed using the described pull-testing setup. For this purpose, the soldering time was varied between two seconds and 120 seconds with five samples per time step. Thermal influence between adjacent solder joint was excluded by cutting the ceramic substrate resulting in single samples with one solder pad each.

### F. Electrical performance

Sheet resistance of the metallization stack during accelerated aging was monitored according to the Van der Pauw method. Cloverleaf structures were integrated in the samples using the described laser process (Figure 2c). The measurement itself was done using a needle prober in combination with a four-point measurement (*Agilent 34401A, Agilent Technologies, Santa Clara, California*). The sheet resistance was calculated with equation (1), while the current *[46] I*_*AB*_ *= 1 mA* was injected whereas the voltage *U*_*CD*_ was measured. Additionally, the electrical conductivity of the solder joints to the thin film layer tracks was determined qualitatively (Figure 2d).

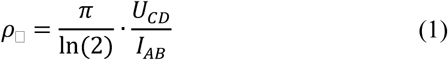

### G. Long-term performance

To evaluate the long-term performance of the material stack the optimized set of processing parameters was used to fabricate substrates for long-term testing (Table 1) containing samples with soldered copper wires and with gold studs.

Accelerated aging was used as a method to simulate the long term behavior and the lifetime of the material stack. For this purpose, the samples were exposed to elevated temperatures in order to accelerate chemical reactions [47]. This method is described in detail in the standard *ASTM F1980 (“Standard Guide for Accelerated Aging of Sterile Barrier Systems for Medical Devices”*) of *ASTM International* (American Society for Testing and Materials)[48]. Based on the Arrhenius equation and the assumption of a constant activation energy *E*_*a*_ of the deteriorating reactions, the time *t*_*0*_ at temperature *T*_*0*_ = 37 °C (for implantable medical devices) can be calculated with the time *t*_*1*_ at the evoked temperature *T*_*1*_ (Equation 2). Herein, the “*Q*_*10*_ *= 2 rule*” is applied, which indicates that a temperature rise of 10 °C causes an increase in the reaction rate by a factor of two [18, 47].

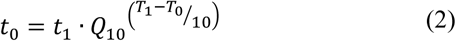

To verify the long term stability of the material stack, different test conditions were applied. First, the samples were encapsulated in medical grade polydimethylsiloxane (PDMS; MED 1000, Nusil Technology, Carpinteria, USA) and exposed to phosphate buffered saline (PBS) at 60 °C to simulate oxidizing and ionic conditions inside the human body. After drying and cooling down to room temperature the PDMS encapsulation was carefully mechanically removed. Due to the mechanical influence the gold studs were no longer representing and therefore not further analyzed in this study. Nevertheless, the adhesive strength of the material stack was determined via pull testing. Appling even higher temperatures for accelerated aging in these settings leads to non-linear side effects as the polymer would alter due to its glass transition temperature [18, 47]. Therefore, to even further increase the aging coefficient, samples were exposed to 150 °C in air as well as in nitrogen atmosphere without PDMS encapsulation. These samples were evaluated on the adhesive strength via pull testing using copper wires and on the stability of the gold studs via shear testing (3.2). Furthermore, leaching of titanium into the platinum metallization can be monitored by a change in sheet resistance of the metal layer [29, 31, 49].

## IV. Results

### A. Initial adhesive strength of different layer thicknesses

The main focus of the present study was the evaluation and optimization of the surface metallization concerning the adhesive strength. The effect of the input values, the prediction functions and its coefficient of determination (r^2^) of the layer stack was considered for the analysis of the DoE. Herein each input parameter combination was measured ten times.

Using tungsten-titanium-alloy (WTi10) as adhesion layer, the main effects corresponded to the following input parameters: 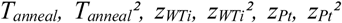 and 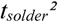, and the interaction of *z*_*Pt*_ with *T*_*anneal*_ as well as *z*_*WTi*_ with *t*_*solder*_. The interaction of *T*_*anneal*_ with *t*_*solder*_ and *t*_*solder*_ itself had minor effects.

A maximum adhesive strength of 62.84 ± 6.55 MPa was achieved with the input parameters *z*_*WTi*_ = 42.9 nm, *z*_*Pt*_ = 100 nm, *T*_*anneal*_ = 101.4 °C und *t*_*solder*_ = 10.38 s. The resulting prediction function (Figure 4, dashed line) showed a coefficient of determination of 0.67. The breakage was of an adhesive nature with cohesive proportions between the solder and the Pt layer for the factor combinations including *T*_*anneal*_ = 20 °C and *T*_*anneal*_ = 360 °C. In contrast, all factor combinations including *T*_*anneal*_ = 700 °C resulted in an adhesive breakage between the ceramic substrate and the metal stack.

**Figure 4:**
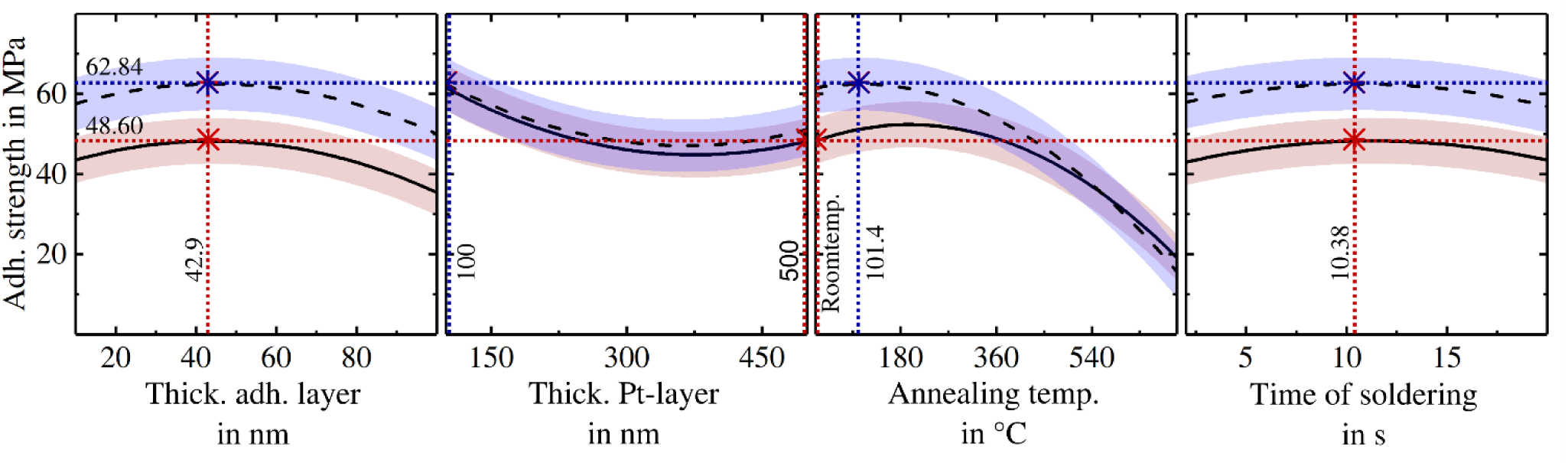
Visualization of the prediction functions of the investigated sputtered layer stacks with tungsten titanium (WTi10) as an adhesion promoter and a surface metallization made of platinum. Dashed/Maximum: The input parameters are optimized for highest adhesive strength. Solid/Optimum: The input parameters are optimized considering the minimization of the specific sheet resistance as well as reducing the complexity of the manufacturing process while keeping the resulting adhesive strength in an acceptable range. Respective values for maximum as well as optimum are given and marked as intersecting lines.

Increasing the thickness of the platinum layer to 500 nm reduced the sheet resistance but decreased the maximum adhesive strength of the layer stack to 51.52 MPa. As there was only a small increase of the breaking strength due to the determined optimum annealing temperature (101.4 °C), the annealing step could be omitted, which resulted in an adhesive strength of 48.60 MPa (Figure 4, solid line).

### B. Layer characterization

The characterization of the metallization layers focused on the surface topography of the platinum metallization and in the validation of the fabrication process. A platinum surface metallization was fabricated using the optimal process parameter discussed in 5.1.1 (Figure 5). The substrate was equally covered with the metallization layer. Additionally, grains became visible which resulted in a surface roughness of *R*_*a*_ *= 449*.*3 ± 28*.*6 nm* with a maximum height of *R*_*t*_ *= 5*.*43 ± 0*.*27 µm* (n = 6, Wyko NT9100, Veeco Instruments Inc., Plainview, NY, USA).

**Figure 5:**
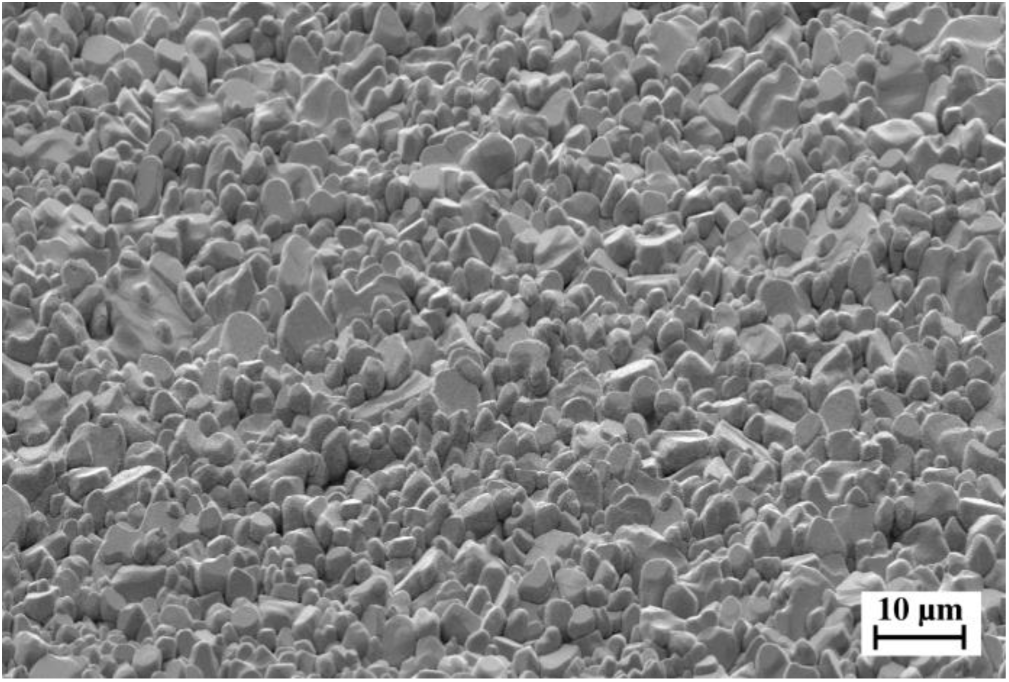
SEM scan of the surface platinum metallization on a ceramic substrate with WTi as adhesion layer with a magnification of 1k. The surface featured a high surface roughness with grains in the range of 2 to 10 µm.

The topography of one single grain got visible, revealing nano-structures, with a magnification of 30k (Figure 6). The cross-section revealed by a FIB of the Al_2_O_3_-WTi-Pt layer stack showed a thickness of the tungsten-titanium layer of 44 nm while the thickness of platinum was approximately 430 nm.

**Figure 6:**
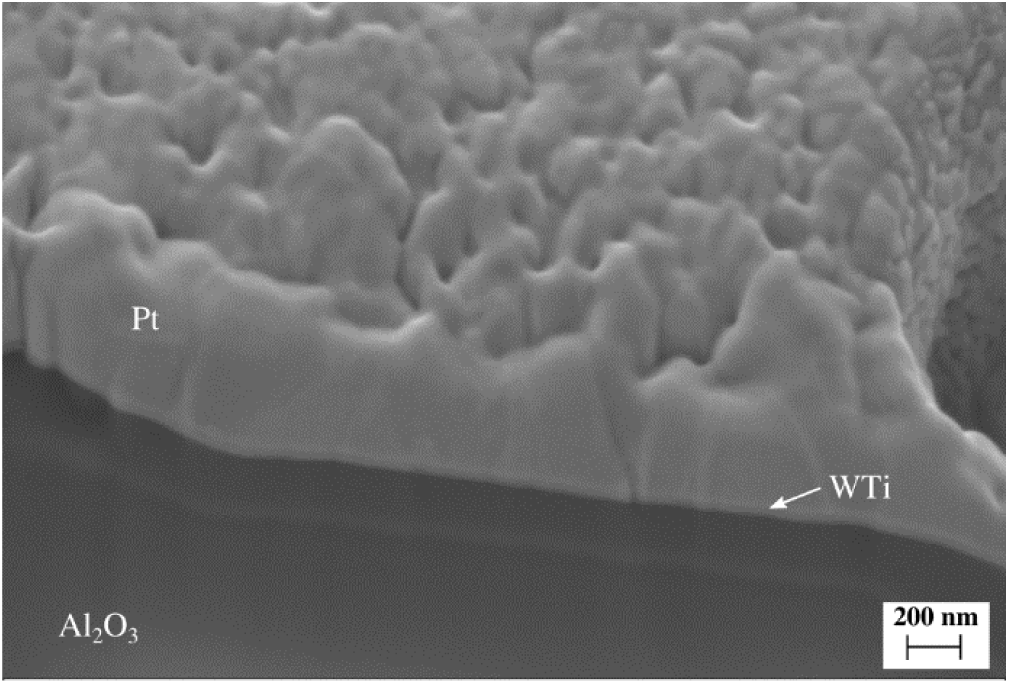
Cross-section of the ceramic, tungsten titanium and platinum layer stack with a magnification of 30k. The thickness of the WTi layer was 45 nm while the platinum layer was 430 nm. In the ceramic a shadowing effect occurred due to electrical charging.

### C. Long-term performance

#### 1) Stability of the assembly with soldered interconnects at 60°C in PBS

To simulate body conditions, samples were encapsulated in PDMS and aged at 60°C for 11 weeks in phosphate-buffered saline, which is equivalent to 54 weeks at body temperature (37°C). Prior to the measurement (pull test), the samples were dried and the PDMS was released. Within the investigated time span, no explicit influence on the adhesive strength could be observed (Figure 7). The adhesive strength resulted in an average of 42.16 ± 7.33 MPa.

**Figure 7:**
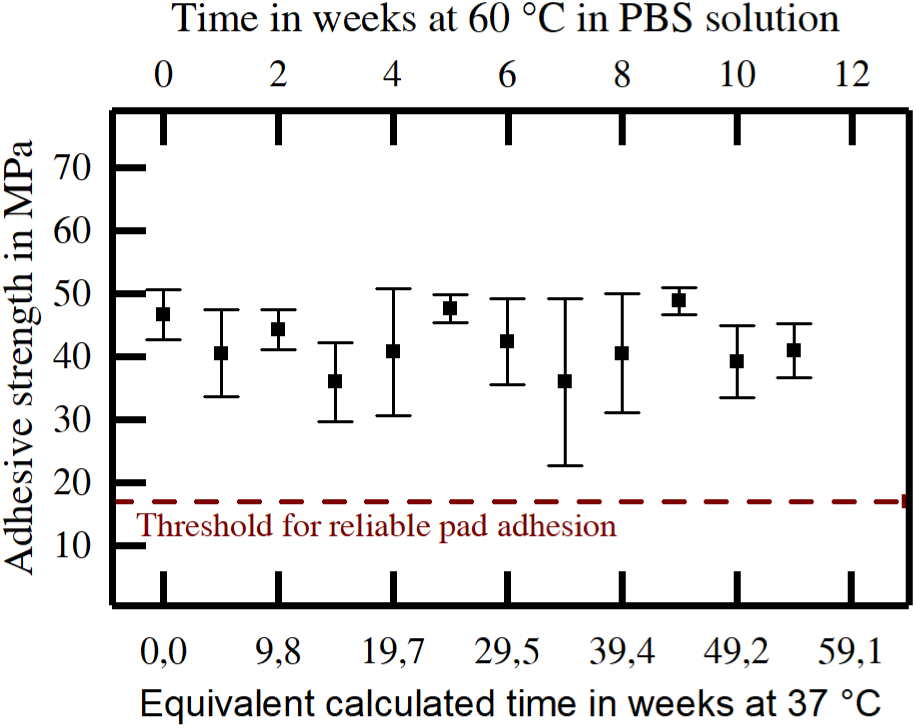
Accelerated aging of PDMS encapsulated samples at 60 °C in phosphate buffered solution. The resulting adhesive strength of the test structures in correlation with the test time and the equivalent calculated time at body temperature is displayed. Devices under test were assemblies with soldered copper wires to the manufactured pads (n=5).

#### 2) Stability of the assembly with soldered interconnects at 150°C in air and nitrogen

The adhesive strength of the samples aged at 150 °C in air decreased from 48.54 ± 7.90 MPa to 32.75 ± 7.08 MPa over the test period of 18 days. Nevertheless, during the entire test period, the adhesive strength was above the threshold for reliable pad adhesion (Figure 8, black data points). The threshold for reliable pad adhesion was set to 17 MPa referring the recommendations for a robust process of comparable screen-printing metallization processes [18]. The test period corresponded (according to equation 1) to a time span of 125 years at body temperature (37°C).

**Figure 8:**
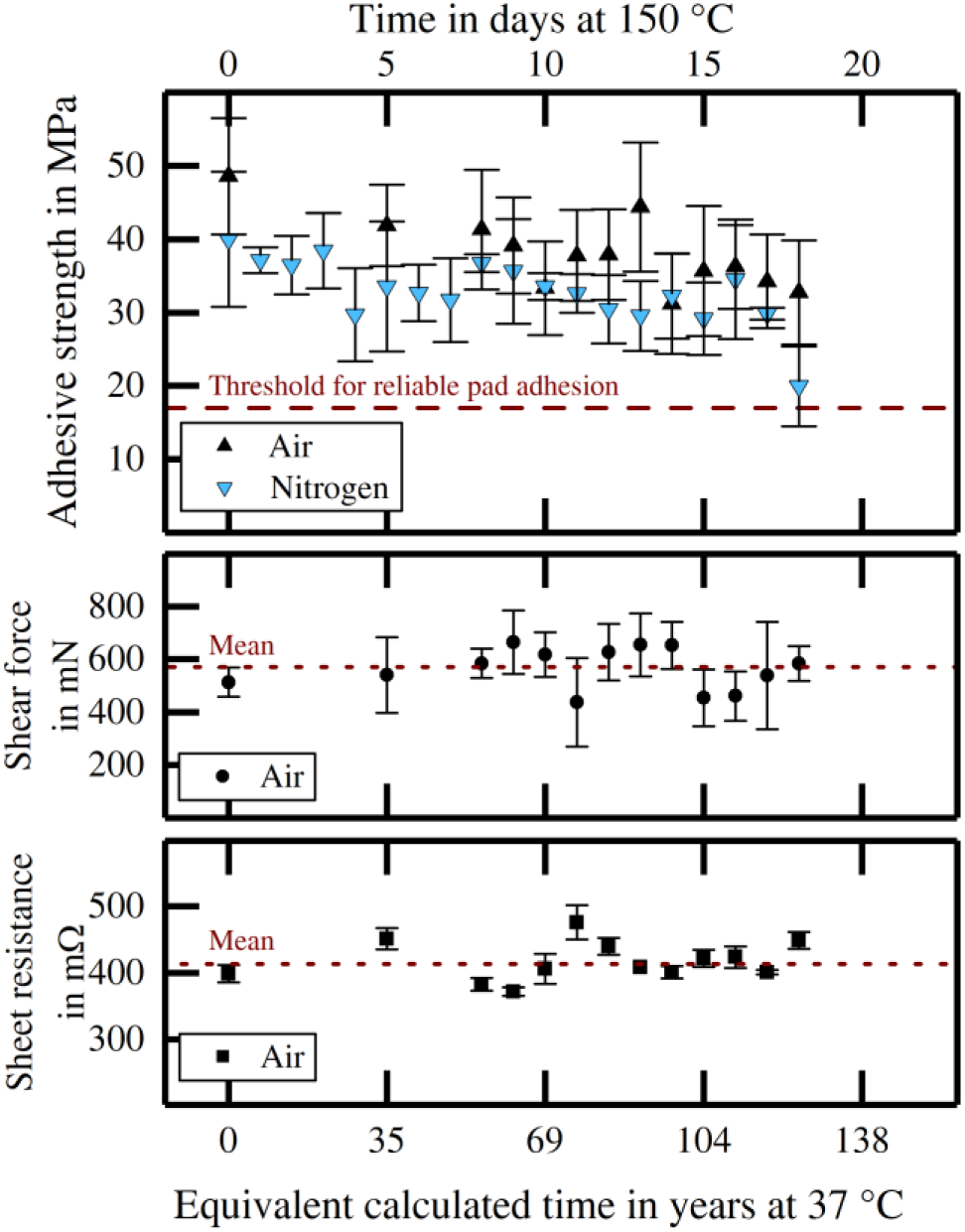
Accelerated aging samples at 150 °C exposed to nitrogen atmosphere (blue) and air (black) during aging. The resulting adhesive strength of the soldered copper wires (top; n=5), the maximal shear force applied to gold studs (middle; n=5) and the sheet resistance of the thin film metallization (bottom; n=2) in correlation with the test time and the equivalent calculated time at body temperature is displayed.

To evaluate the influence of oxygen in the atmosphere, the accelerated aging was repeated in nitrogen atmosphere at 150 °C for 18 days. It was observed, that there were comparable changes in the adhesive strength over the entire test period as observed in air (Figure 8, blue data points). The adhesive strength dropped from 39.96 ± 9.18 MPa (day 0) to 29.81 ± 0.80 MPa (day 17). On day 18, a high change in the adhesive strength was observed that resulted in a mean value of 19.93 ± 5.44 MPa (Figure 8).

An exemplary EDX measurement was executed after pull-testing and accelerated aging in nitrogen atmosphere at 150°C. The measurements analyzed the left-over breaking edge on the ceramic substrate after pulling the copper wire. Herein, the mole fraction of samples in different aging stages was measured. The main changes, between day 0 and day 18, were an increase in the mole fraction of oxygen (*x*_*O,0*_ = 20.7 ± 2.7 % to *x*_*O,18*_ = 53.2 ± 0.4 %) and aluminum (*x*_*Al,0*_ = 0.0 % to *x*_*Al,18*_ = 33.4 ± 0.7 %), while platinum (*x*_*Pt,0*_ = 11.9 ± 2.5 % to *x*_*Pt,18*_ = 1.16 ± 0.1 %), tin (*x*_*Sn,0*_ = 44.8 ± 4.0 % to *x*_*Sn,18*_ = 5.8 ± 0.6 %) and lead (*x*_*Pb,0*_ = 22.2 ± 0.3 % to *x*_*Pb,18*_ = 3.7 ± 0.1 %) decreased. The mole fraction of titanium (*x*_*Ti,0*_ = 0.0 % to *x*_*O,18*_ = 1.1 ± 0.1 %) and tungsten (*x*_*W,0*_ = 0.0 % to *x*_*O,18*_ = 1.7 ± 0.1 %) showed minor changes.

#### 3) Stability of the assembly with wire-bonded gold studs

The maximum shear force applied to the bond studs showed no clear trend during accelerated aging for 18 days in air at 150 °C. An average over the investigated time of the shear force of 570.27 ± 128.34 mN was obtained (Figure 8). Optical evaluation of the breakage revealed a cohesive breakage within the gold phase of the bond studs at every time over the 18 days.

#### 4) Sheet Resistance of the metallization at 150°C in air

The sheet resistance of the metallization was monitored in an interval of 18 days during accelerated aging at 150 °C in air, which is (according to equation 1) equivalent to 125 years at body temperature (37 °C). Within the investigated time span, the measurement showed no crucial influence. The sheet resistance was measured at an average of 413.5 ± 27.5 mΩ (Figure 8). The qualitative investigation of the electrical conductance of the solder joints to the thin film metallization showed no influence during accelerated aging.

#### 5) Influence of soldering time

The time of soldering showed low influence to the resulting adhesive strength, varying between 34.28 ± 3.54 MPa at t_solder_ = 2 s and 47.33 ± 16.12 MPa at t_solder_ = 120 s (Figure 9). According to the mean values, a minor positive trend can be observed, which cannot be specified due to the large variance in the data. Nevertheless, the resulting adhesive strength was above the threshold for reliable pad adhesion at all time.

**Figure 9:**
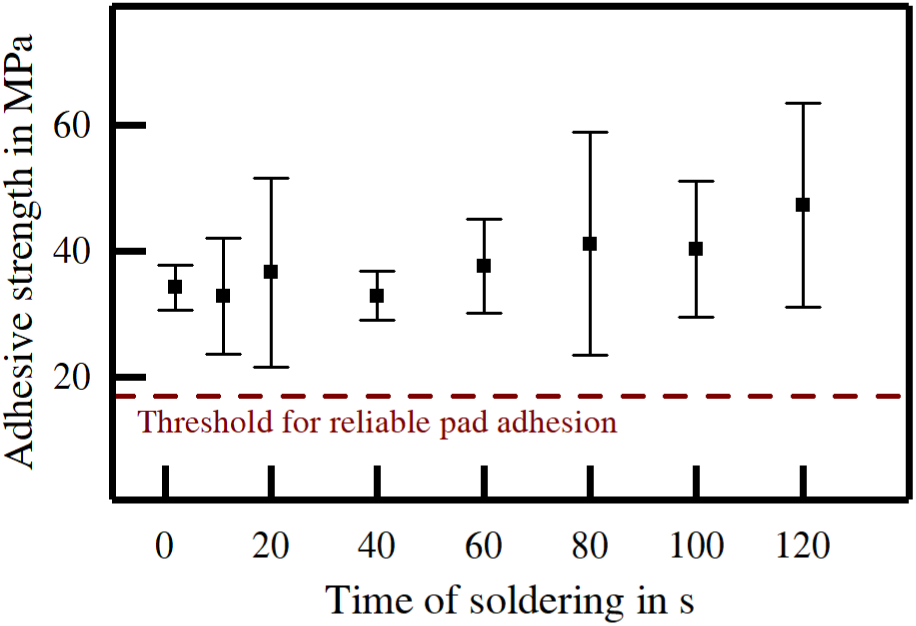
Influence of t_solder_ on the adhesive strength during soldering (n=5).

## V Discussion

The performed DoE with 16 different combinations of processing parameters enabled the generation of a mathematical prediction model for the adhesion of a material stack consisting of alumina substrate, titanium adhesion promoter, functional platinum layer and soft soldering of interconnects. The obtained model allowed an optimization of the material stack concerning its mechanical performance. Furthermore, investigations on the long-term behavior of the assembly showed stable operation of soldered connections as well as gold stud interconnects applied by wire bonding.

### A. Initial characteristics of the manufactured metallization

#### 1) ptimal manufacturing parameters

Increasing the thickness of the platinum surface metallization from 100 nm to 500 nm leads to a drop in the adhesive strength of the metallization stack. However, the resulting adhesive strength is still almost three times the threshold for reliable pad adhesion (17 MPa) while the electrical conductivity is substantially improved. Omitting the annealing step (*T*_*anneal*_) simplifies the manufacturing process with only minor losses in the initial mechanical properties and prevents leaching of titanium into platinum as can be observed at temperatures above 270°C. Furthermore, the wide range and curve shape of *T*_*anneal*_ could indicate a higher order correlation. Therefore, we set the optimal input parameters for the fabrication of thin film platinum metallization on alumina substrate to 43 nm for the thickness of the adhesion promoter (WTi10) and 500 nm for the thickness of the platinum surface metallization with no dedicated annealing step. Additionally, there was no strong dependency of *t*_*solder*_ on the adhesive strength of the material stack with soldered copper wires. Thus, an influence of the individual soldering skills on the metallization (i.e. the time needed for applying a proper solder joint) can be neglected.

#### 2) Electrical properties at optimal manufacturing parameters

The specific resistance of sputter deposited platinum on sputter deposited titanium measured in average 0.207 ± 0.014 µΩm when a metallization thickness of 500 nm is assumed. This is approximately double the bulk specific resistance of platinum, which is in accordance with simulations of electrical conduction in thin metallic films[50] and comparable to reported values of 0.145 µΩm[29] and 0.162 µΩm [31].

Supported by the obtained data and the developed model it can be stated, that the material stack is a valid method to form electrical conductors and interconnection pads on alumina substrates with adhesive properties above critical thresholds for reliable pad adhesion.

### B. Layer characterization

The surface topography of the metallization featured excellent properties for further assembly techniques like soldering or bonding. Due to the roughness, the effective surface was drastically enlarged providing more possibilities for chemical and physical bonding mechanisms. The structures occurred based on two different reasons. First, the grains in the µm range reflected the topography of the underlying ceramics (Figure 5). The average surface roughness of metallized and bare substrates differed about Δ*R*_*a*_ *= 30 nm* or Δ*Rt = 1*.*56 µm*, whereas the roughness of the bare ceramics was slightly higher. Second, the nano-roughness (Figure 6) occurred due to the layer growth of the metallization during sputtering [34]. The cross section of the layer stack revealed a uniform adhesion layer and clear phase boundaries between the different materials, which provided continuous bonding mechanism from the ceramics to the adhesion layer and further to the platinum layer. The thicknesses of the metallization matched with the set parameters. Due to the nano-roughness of the surface this value couldn’t be determined precisely. In Figure 6 an additional layer seemed to appear in the ceramic substrate. In fact, this layer was caused by charging effects of the non-conductive ceramic during the SEM scan and did not exist in reality.

### C. Long-term performance

The long-term performance of the material stack with optimal manufacturing parameters was assessed by evaluating the transient behavior of the sheet resistance, the adhesive strength of soldered copper wires and the maximum shear force of gold studs. Supported by x-ray spectroscopy measurements a conclusive picture of the long-term behavior of the material stack was created.

#### 1) Sheet Resistance

Previous studies showed substantial increase in specific resistance when titanium is leaching into platinum forming intermetallic phases, oxides and microholes [29, 31]. In the here presented work no changes in the specific resistance were observable during accelerated aging for 18 days at 150 °C in dry conditions. Therefore, it is not likely that an excessive alteration of the manufactured metallization layers took place under these conditions. Additionally, the presence of solder showed no influence on the electrical behavior during these 18 days. In terms of electrical properties, the present thin film metallization showed excellent preconditions for implantable applications where temperatures are as low as 37 °C. Nevertheless, the metallization still has to be investigated under wet conditions, including the desired encapsulation.

#### 2) Adhesive strength

The results of the accelerated aging studies (150 °C in air/nitrogen and 60 °C in PBS solution) showed that the present thin film metallization features an adhesive strength of at least 17 MPa within extrapolated 127 years in air/nitrogen, respectively 54 weeks in solution. These values are above the threshold for reliable pad adhesion for robust processes, which is set to 17 MPa. In consequence of a lack of an appropriate threshold for thin film metallization, this value is adopted from the manufacture recommendation for the adhesive strength of screen printing processes used for the manufacturing of medical implanted PCBs [18]. The breakage occurred within the metallic phase (solder, platinum, titanium) of the layer stack in each case. This breakage is either of cohesive or adhesive nature. Nevertheless, no adhesive failure of the metal ceramic interface was observed. Thus, the presented values do not represent the actual adhesive strength of the ceramic metallization interface. In fact, the results suggest that the critical ceramic metal interface features at least the presented adhesive strength [16]. Hence, the interface shows excellent properties.

Regarding the adhesive strength of the soldered copper wires, a slight loss in the adhesive strength can be observed over time in air and nitrogen at 150°C. The measurements of the mole fractions of the break edge on the substrate show an increase of aluminum and oxygen, and were accompanied by a decrease of platinum, tin and lead. This behavior may be an indicator for diffusion of the thin film layers into the overlying solder [28] or for a loss of integrity of the titanium layer [31]. In both cases the breakage point would get closer to the ceramics, whereby the interaction volume of the EDX measurement penetrates more into the substrate. This corresponds to the increase in aluminum and oxide in the break edge. The observed decrease in adhesive strength with an increase in *T*_*anneal*_ during the preliminary investigations also renders both statements possible, since diffusion in solid materials is a process accelerated by elevated temperatures. It must be noted here that the test temperature (150 °C) is very close to the melting temperature of the solder (179 °C). Diffusion processes in the solder phase could thus be further affected. However, there should be no influence on the metal-ceramic interface, especially since the weakest point is always in the metallic phase, as mentioned earlier.

It has previously been shown that the interface gold bond/sputter deposited platinum does not get visually altered when aging up to the recrystallization temperature of gold (225 °C) in air [36]. Likewise, in the presented work no trend in maximum shear force of gold stats was visible during 18 days in air at 150°C. This trend can also be observed in the presented work. However, the breakage after shear testing was of cohesive nature in the gold phase at all times. Therefore, a loss in adhesive strength could be present, but out of the capability of the measurement method when the gold stud is collapsing before the underlying material stack.

Nevertheless, the fact that the breakage of the bond studs was always in the gold phase shows the potential of the present thin film metallization for gold bond interconnection in long term applications without any change in the mechanical properties.

In conclusion it can be said that all observations support the usability of the investigated technology to form high reliable and long term stable electrical conductors and interconnects in demanding low-temperature applications like medical implants.

#### 3) Usability for fabricating medical implants

Liquidus diffusion can occur during assembly of soldered parts and is a challenge in other process technologies in medical implants [18]. Regarding implantable systems, soldered parts are not only limited to electronics, but can as well be e.g. hermetic sealings [51]. However, for long term application, it is important to guarantee a well fabricated solder joint, which might require some time during soldering, especially with manually conducted fabrication steps. It seems reasonable to consider that soldering times up to two minutes are needed for these processes. The presented data shows that during soldering within this time span the adhesive strength of the present thin film metallization is not altered. Therefore, it is safe to assume that a liquidus diffusion might occur in the metallic phase [28], but the adhesive bonds between the metallization layer stack and the ceramic substrate will not mutate. In contrast, with state-of-the art metallization consisting of platinum-gold metallization embedded in a glass matrix, liquidus diffusion has a higher impact [18]. In this case a soldering time exceeding approximately 20 seconds leads to catastrophic failures of the system [18]. Therefore, it is advantageous to replace previously used glass-based planar metallization with the sputter-deposited material stack described in this work.

## VI. Conclusion

In the presented work, a new material combination for metalizing alumina substrates is thoroughly investigated adducing crucial *in vitro* test scenarios for the application in the field of AIMDs. In particular, the focus was on the usage of this metallization for both hermetic and non-hermetic approaches. In both applications the entire system is coated by a protective medical grade silicone rubber layer, limiting diffusion of ions and making tests on cellular or systemic toxicity of the metallization layer obsolete, as demonstrated by similar chronic implantation studies [52, 53]. Instead, the electrical and mechanical long term performance, especially in combination with commonly used assembly methods, is of high interest in the manufacturing of such AIMDs.

The developed sputter deposited WTi-Pt metallization showed excellent stability over long durations at elevated temperatures in air and in saline solution. The stability to soft solder as well as to gold-stud bonds on this metallization epitomized its significance for medical implant manufacturing. All manufacturing steps were conducted at low temperatures compared to state of the art ceramic metallization techniques which removed any restrictions that CTE-mismatch or pressure-distributions during high temperature firing pose on the shape of the ceramic to be metallized. Therefore, the presented process allowed for heterogeneous assemblies on ceramic substrates with degrees of freedom and longevity that had not been reported before.

The thin film process enables reliable electrical and mechanical connection by riveting multi-channel arrays via with gold-studs [41, 54] and by soft soldering any kind of wire [18]. Regarding the latter, one valuable result of the presented studies is the outstanding stability of the material stack during soft soldering. These findings, in combination with the described mask-less structuring process, render the developed thin film metallization to be a promising technology for hermetic and non-hermetic active implantable medical devices. The stable process and structural accuracy with the excellent long-term stable properties opens a wide application field for the presented technique, including fine-pitch highly stable conductors and interconnection pads for implants [55].

Based on the ongoing investigations of the presented study the material stack has been reported to successfully seal hermetic metal/ceramic enclosures using soft solder as well as hermetic electrical feedthrough channels applying gold-studs. [56]

## Acknowledgment

The authors would like to thank the KNMF at Karlsruhe Institute of Technology for the granted working hours at the FIB (ID 2017-018-019571).

